# Epileptiform GluN2B–driven excitation in hippocampus as a therapeutic target against temporal lobe epilepsy

**DOI:** 10.1101/2021.06.30.450508

**Authors:** Adam Gorlewicz, Barbara Pijet, Krstina Orlova, Leszek Kaczmarek, Ewelina Knapska

## Abstract

NMDAR is an ionotropic glutamate receptor critically involved in excitatory synaptic transmission. The receptor properties are strongly determined by its subunit composition. One of the NMDAR subunits is GluN2B, which displays restricted and spatially different from other subunits expression in the mature brain. GluN2B–containing NMDARs are present in the hippocampus – a structure playing a major role in temporal lobe epilepsy (TLE). However, the contribution of GluN2B to pathophysiology of TLE has not been fully explored. Here, we report the functional alterations of GluN2B–containing NMDAR receptors in the hippocampus in distinct mouse models of temporal lobe epilepsy. In particular, we show the impact of GluN2B on excitatory feedback in granule cells. Based on these results, we propose a mechanism–oriented effective antiepileptic strategy that selectively antagonizes GluN2B–containing NMDARs with ifenprodil, a well–known GluN2B antagonist. Collectively, our research identifies GluN2B as one of the pivotal factors in pathogenesis of temporal lobe epilepsy and associated recurrent seizures. Furthermore, our study indicates the prospective antiepileptic properties of ifenprodil in TLE.

## Introduction

Temporal lobe epilepsy (TLE) is the most common form of human epilepsy and is characterized by recurrent partial seizures originating from such mesial structures as the hippocampus, propagating to the other limbic constituents, and finally giving rise to general convulsions (Bertram, 2009). Over the last decades, a new generation of antiepileptic drugs approved for treatment of TLE has emerged. However, satisfactory control of chronic seizures is still missing for about 40% of patients for whom the disorder is refractory to currently available pharmacotherapies (de Lanerolle and Lee, 2005). For this reason, TLE is still a serious problem and new therapeutic strategies are desired.

In TLE, epileptogenesis is a progressive cascade of molecular and cellular modifications. Over the course of these modifications, neurons undergo reconfiguration associated with alteration of their connectivity and excitability, and gradually evolve into a seizure–prone hippocampal network (Pitkänen and Lukasiuk, 2011). Epileptogenesis extends beyond clinical TLE with associated recurrent seizures, contributing not only to the development, but also to the progression of the epileptic condition once it is established (Pitkänen et al., 2015). Hence, identification of the basic operative mechanisms of epileptogenesis might guide treatments that could ameliorate the condition. Many studies on the origin of TLE concentrate on a synaptic pathology in which the NMDAR–dependent excitatory glutamatergic system plays a key role (Masukawa et al., 1991; Rice and DeLorenzo, 1998; Abegg et al., 2004). For instance, molecular and microscopic investigations demonstrate alterations in NMDAR_GluN2B_ neuronal membrane localization and cellular expression. Changes such as these are also expected according to the epileptogenesis induced by self–sustaining status epilepticus (SSSE) model (Frasca et al., 2011). Experiments carried out *ex vivo* on organotypic hippocampal cultures have shown that selective inhibition of NMDAR_GluN2B_ reduced both the epileptiform rearrangement of the hippocampal network and epileptic discharges of dentate gyrus (DG) granule cells (GC) (Wang and Bausch, 2004). This suggests that GluN2B might play an essential role in epileptogenesis, and that selective antagonism of NMDAR_GluN2B_ might be a fruitful way to treat TLE. Nevertheless, this hypothesis has never been sufficiently tested *in vivo*.

Ictogenesis is a process of seizure generation by an epileptiform network (Blauwblomme et al., 2014). In developed TLE, the hippocampus has a constant susceptibility to seizures (ictal activity) (Alexander et al., 2016) separated by much longer interictal intervals. While our understanding of the primary mechanisms responsible for the network transition from the interictal period to seizures is not complete. At a very basic level it results from discrete, progressive changes in neuronal population dynamics corresponding to a reduction of network homeostasis, and as a result, to abnormally synchronous neuronal activity. Ictogenic processes are one of the prime targets in medication aiming at suppression of seizure initiation (Blauwblomme et al., 2014; Lillis and Staley, 2018).

Because the classic view on loss of homeostasis in seizure in TLE implicates imbalance between excitation and inhibition, glutamatergic transmission is considered to play a critical role in ictogenesis. Among glutamate receptors, a substantial amount of data points to NMDAR as an important factor in the pathophysiology of seizure initiation (Hanada, 2020). While the role of NMDAR_GluN2B_ in the origin of seizures has been well examined in models of acute convulsions (Pontecorvo et al., 1991; De Sarro et al., 1995, 1997; Doyle and Shaw, 1996; Tsuda et al., 1997), it has not been sufficiently studied in recurrent seizures in TLE.

Ifenprodil, a selective antagonist of NMDAR_GluN2B_, has been demonstrated to decrease neuronal excitability in lateral temporal lobe and neocortex resections from epileptic patients with cortical dysplasia (Möddel et al., 2005; Wang et al., 2017). Moreover, a combination of NBQX and ifenprodil, two distinct antagonsists acting at different glutamate receptor subtypes, deploy antiepileptogenic effects in the kainate model of TLE (Schidlitzki et al. 2017). Despite that, the direct effect of ifenprodil on epileptogenesis and chronic seizures in TLE remains unknown. Our results reported herein and obtained in rodent TLE models suggest that a specialized subpopulation of N– metylo–D–aspartate receptor (NMDAR) containing the GluN2B subunit (NMDAR_GluN2B_) accounts for a substantial component of aberrant synaptic transmission in epileptiform hippocampal networks. This opens a new perspective for an effective epilepsy treatment through targeting of NMDAR_GluN2B_ during epileptogenesis and ictogenesis. Ifenprodil offers an interesting therapeutic option in this regard.

## Material and methods

### Animals

Mice were kept in the animal house of the Nencki Institute. Some of the experiments were performed on mice with a GluN2B gene flanked by loxP sites (GluN2B^ff^) –a generous gift of Laboratory of Synaptic Circuits of Memory (IINS Bordeaux). All animals were housed in cages with 12h light / dark schedules. Water and food was available *ad libitum*. All the experiments on animals were conducted based on national laws that are in full agreement with the European Union directive on animal experimentation (Directive 2010/63/EU).

### Generation of conditional GluN2B^−/−^ mice

GluN2B^ff^ mice were generated as described by von Engelhardt (2008) (von Engelhardt et al., 2008). To produce DG selective GluN2B^−/−^ mice, we used a lentiviral vector expressing Cre– recombinase under control of C1ql2 minimal promoter. The vector was produced by the viral vectors facility of Bordeaux “TransBioMed”. In a typical procedure, male GluN2B^ff^ animals (P40 – P60) were anesthetized by isoflurane inhalation, placed in a stereotactic frame, and injected with buprenorphine to prevent post–surgery pain. After that, 100□nL of viral solution (10^9^ particles) were injected intra– cranially in a localized fashion using a micropump and syringe (Nanofil WPI) at a rate of 100□nl / min into the left DG (AP −2.00 mm, ML 1.50 mm, DV −1.90 mm, relative to bregma). After injection, the capillary was maintained *in situ* for an additional 2 min to limit reflux along the injection track. The craniotomy was sealed with silicon elastomer and the scalp was closed with sutures.

### Pilocarpine model of chronic epilepsy

Male FVB mice (P40 – P60) were injected intraperitoneally with a single dose of methylscopolamine nitrate (1 mg / kg) (Sigma–Aldrich) (Turski et al., 1984) to block peripheral effects of pilocarpine. Twenty minutes after the methylscopolamine nitrate administration, a single dose of pilocarpine hydrochloride (Sigma–Aldrich) was injected intraperitoneally (265 mg / kg). Control animals were injected with 0.9 % NaCl. After that, animals were continuously observed for 2 hours. Typically within 45 minutes after the pilocarpine administration the animals experienced convulsions that were classified using the Racine scale (Racine, 1972). Only animals with continuous tonic–clonic (grand mal) seizures were subjected to further experiments.

### Kainate model of chronic epilepsy

Male C57Bl6, GluN2B^ff^, GluN2B^GC+/+^, and GluN2B^GC−/−^ mice (P40 – P60) were stereotactically injected with 50 nl of 20 mM solution of kainic acid (Tocris / Biotechne) in 0.9 % NaCl to the left CA1 area of the dorsal hippocampus (AP −1.8 mm, ML = −1.7 mm, DV −2.1 mm, relative to bregma) under isoflurane anaesthesia (induction 5%, maintenance 2 – 3 %). Control animals were injected with 0.9 % NaCl. Injections were done through a thin glass capillary tube of 50 – 100 μm diameter at the tip using a Hamilton syringe. After injection, the capillary was maintained *in situ* for an additional 2 minutes to limit a reflux along the injection track. The craniotomy was sealed with silicon elastomer, and the scalp was closed with sutures. Directly after the surgery, animals received subcutaneous injection of Carprofen (5 mg/kg).

### Immunofluorescence

Perfusion–fixed (4 % PFA in PBS) brains were dissected and post–fixed by immersion in 4 % paraformaldehyde solution in PBS, pH 7.4 (overnight at 4 °C), then cryoprotected in 30 % sucrose in PBS (up to 48 h at 4 °C), and finally frozen in isopentane cooled in a liquid nitrogen. All specimens were stored at −70 °C until use. Free–floating slices of cryosectioned (50 μm–thick) brains were subjected to antigen retrieval for 10 minutes at 37 °C using Pepsin Reagent (Abcam). This step was followed by 30–minute incubation with aqueous solutions of 5 % normal donkey serum (NDS) (Jackson Laboratories) and 0.2 % Triton X–100 (Sigma) at room temperature (RT), followed by overnight incubation with primary antibody diluted in 5 % NDS with 0.2 % Triton at 4 °C. The detection of immunoreactions was performed using species–specific secondary antibodies (two–hour incubation at RT) that were diluted in 5 % NDS with 0.5 % Triton. After incubation with secondary antibodies, samples were rinsed with PBS, co–stained with DAPI, dried on coverslips, immersed in Vectashield (Vector Laboratories), and finally coverslipped. The following primary antibodies and their dilutions were used: rabbit anti GluN2B (Frontier Institute, 1:100) and rabbit anti synaptoporin (Synaptic Systems, 1:400). The following secondary antibodies and their dilutions were used: donkey anti rabbit conjugated to Alexa488 (Invitrogen, 1:200) and donkey anti rabbit conjugated to Alexa555 (Invitrogen, 1:200).

Fluorescent specimens were examined under a spectral confocal microscope (Leica TCS SP8 with the following objective lenses: HCX PL APO CS 10x / 0.40 DRY, HC PL APO CS2 20x / 0.75 Imm, HC PL APO CS2 40x / 1.30 Oil, HC PL APO CS2 63x / 1.20 Water, HC PL APO CS2 63x / 1.40 Oil) using 488 nm Ar, 594 nm HeNe, and 405 nm diode laser lines for the excitation of Alexa488, Alexa555 and DAPI respectively. To avoid cross–reactivity of the detection systems, the spectral ranges of detectors were adjusted and images were scanned sequentially.

### Patch–clamp in brain sections

Parasagittal hippocampal slices (350 μm–thick) were obtained from C57BL6 mice by cutting with a VT 1200 Vibratome (Leica) in an ice–cold solution containing the following (in mM): 80 NaCl, 2.5 KCl, 25 NaHCO_3_, 1.5 NaH_2_PO_4_, 7 MgCl_2_, 0.5 CaCl_2_, 10 glucose, and 75 sucrose and equilibrated with 95 % O_2_ and 5 % CO_2_. The slices were then incubated at 33 °C for 30 min and subsequently stored at RT in an extracellular medium composed of (in mM): 125 NaCl, 2.5 KCl, 1.25 NaH_2_PO_4_, 25 NaHCO_3_, 2 CaCl_2_, 1 MgCl_2_, 11 glucose (equilibrated with 95 % O_2_ and 5 % CO_2_). For recordings, slices were transferred to the recording chamber in which they were continuously superfused with extracellular medium composed and equilibrated as above.

Whole–cell voltage–clamp recordings were made at 32 °C from DG granule cells under infrared differential interference contrast imaging using borosilicate glass capillaries (resistances between 3.5 – 4 MΩ) filled with a solution containing (mM): 120 cesium methanesulfonate, 2 MgCl2, 4 NaCl, 5 phospho–creatine, 2 Na2ATP, 20 BAPTA, 10 HEPES, 0.33 GTP. The solution was adjusted to pH 7.3 using CsOH. To record NMDAR–mediated currents NBQX (20 μM), bicuculline (10 μM), and CGP55845 (3 μM) were added to the extracellular medium, the membrane potential was clamped at +40 mV, and NMDAR_GluN2B_ channels were antagonised with the presence of ifenprodil (1 μM) in extracellular fluid. Only neurons with resting membrane potential more negative than −55 mV were used. Access resistance was monitored during the entire recording time with application of a hyperpolarizing voltage step (−5 mV, 10 ms) once every minute, and when it changed by >20 %, the recording was excluded. Whole–cell patch clamp recordings were made using Multiclamp 700B amplifier (Molecular Devices) connected to the Digidata 1440B system (Molecular Devices). Data were acquired with pClamp (Molecular Devices), digitized at 20 kHz, and filtered at 3 kHz.

### Extracellular recordings in brain sections

Parasagittal hippocampal slices (450 μm–thick) were obtained from control and kainate– treated C57BL6 mice by cutting with a VT 1200 Vibratome (Leica) in an ice–cold solution containing the following (in mM): 80 NaCl, 2.5 KCl, 25 NaHCO_3_, 1.5 NaH_2_PO_4_, 7 MgCl_2_, 0.5 CaCl_2_, 10 glucose, 75 sucrose and equilibrated with 95 % O_2_ and 5 % CO_2_. Afterwards, sections were transferred to the interface chamber, placed on nylon mesh, and kept at ~34 °C in the interface between warm, humidified carbogen gas (95 % O_2_, 5 % CO_2_) and extracellular solution (pH 7.4) containing (mM): 125 NaCl, 7.5 KCl, 2.3 CaCl_2_, 1.3 MgCl_2_, 1.25 NaH_2_PO_4_, 26 NaHCO_3_, 20 glucose. Sections remained in extracellular solution for 40 minutes. After that time the medium was exchanged with another extracellular solution (pH 7.4) containing (mM): 125 NaCl, 7.5 KCl, 2.3 CaCl_2_, 0 MgCl_2_, 1.25 NaH_2_PO_4_, 26 NaHCO_3_, 20 glucose. Changes of local field potentials (LFP) were recorded in the DG with a sixteen–shank (50 μm–thick) silicone electrode with 200 μm separation distance between the shanks (Atlas Neuroimaging) or with silver wire single macroelectrode (250 μm–thick). Multichannel recordings were amplified using an Intan chipset (RHD2132, Intan Technologies) connected to an Open Ephys acquisition board (Open Ephys). Multichannel data were digitalized at 10 kHz and filtered at 3kHz. Single channel recordings were made using a DP-311 differential amplifier (Warner Instruments) connected to the Digidata 1440B system (Molecular Devices). Data were acquired with pClamp (Molecular Devices), digitized at 20 kHz, and filtered at 3 kHz.

### EEG electrode implantation and signal acquisition

C57BL6 mice one month after the either kainate or NaCl treatment were subjected to the isoflurane anaesthesia (induction 5 %, maintenance 2-3 %). The scalp was removed and a bipolar hippocampal electrode (Bilaney Consultants GmbH) was implanted into the left hippocampus (AP - 2.0 mm, ML 1.3 mm, DV −1.7 mm, relative to bregma). In addition, four stainless–steel screw electrodes (1.6 mm ∅, Bilaney Consultants GmbH) were implanted. One screw electrode was placed ipsilaterally to the hippocampal craniectomy over the frontal cortex and another one contralaterally to the craniectomy also over the frontal cortex. A reference screw electrode and a ground screw electrode were positioned over the cerebellum (see Fig. 4). Prior to the video EEG monitoring (vEEG), mice were placed in Plexiglas cages (one mouse per cage) and connected to the recording system with commutators (SL6C, Plastics One). vEEG was performed using the TWin EEG recording system that was connected to a Comet EEG PLUS with AS40 PLUS 57–channel amplifier (Grass Technologies) and the signal was filtered (high–pass filter cut–off 0.3 Hz, low–pass filter cut–off 100 Hz, 50 Hz noise eliminator). The video was recorded using an I–PRO WV SC385 digital camera (Panasonic).

**Fig 1.**
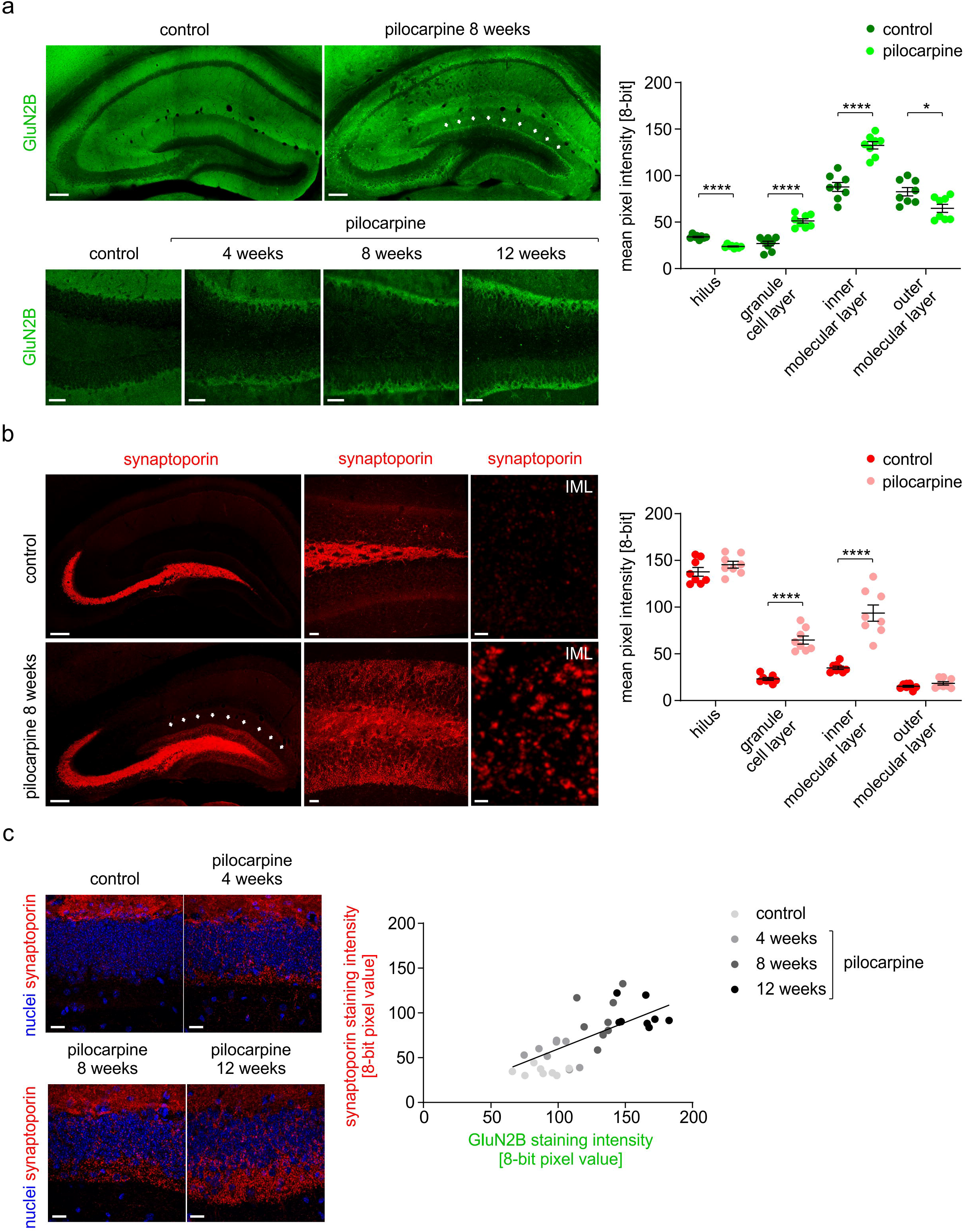
GluN2B in hippocampal network rearrangement during epileptogenesis. **a)** NMDAR_GluN2B_ immunoreactivity is increased in the IML of pilocarpine–treated chronic animals when compared to control specimens (arrows in top right panel). Increased NMDAR_GluN2B_ immunoreactivity can be detected at different stages of TLE development, up to at least 12 weeks following convulsant administration (bottom panels). Mean fluorescence intensity analysis in different subregions of the DG quantitatively demonstrates NMDAR_GluN2B_ immunoreactivity alteration in the DG of pilocarpine–treated animals (n = 8) in comparison to control mice (n = 8) (graph – p < 0.0001 for hillus, GCL, IML and p = 0.013 for OML, t–test). Scale bars: top panels 250 μm, bottom panels 70 μm **b)** Immunoreactivity of synaptoporin in control (top panels) and pilocarpine–treated epileptic animals (bottom panels). Increased expression of synaptoporin in GCL and IML (top panels – arrows) of TLE animals in comparison to control conditions indicates the presence of mossy fiber sprouting as a consequence of neuronal network alteration in the epileptic hippocampus. Mean fluorescence intensity analysis in different subregions of the DG quantitatively demonstrates TLE–related synaptoporin immunoreactivity alteration in the DG (n = 8 for both investigated groups) (graph – p < 0.0001 for GCL and IML, p = 0.21 for hilus, and p = 0.15 for OML, t–test). Scale bars: left panels 250 μm, middle panels 20 μm, right panels 3 μm **c)** Altered NMDAR_GluN2B_ expression in the IML corresponds to the localization of sprouted mossy fiber terminals in that region as indicated with synaptoporin immunoreactivity (top panels). Mean fluorescence intensities of both immunoreactivities in that region increase in time as TLE develops which implicates correlation between NMDAR_GluN2B_ and sprouted mossy fiber terminals (graph – r = 0.74). Scale bars: 20 μm

**Fig 2.**
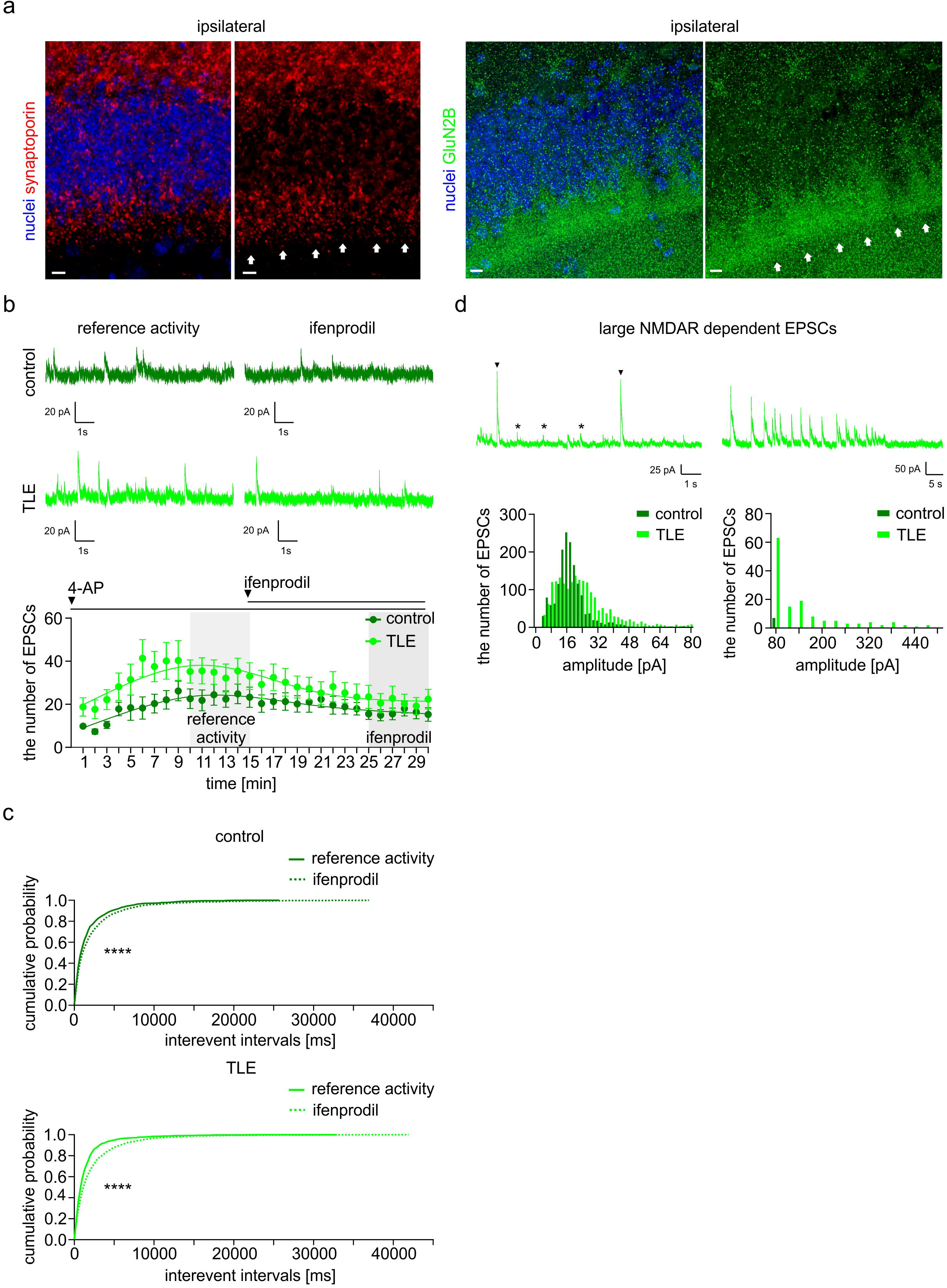
Altered NMDAR_GluN2B_–dependent synaptic transmission at epileptiform granule cells of dentate gyrus. **a)** Immunoreactivity of synaptoporin in the DG of kainate–treated animals demonstrating the presence of synaptoporin–positive sprouted mossy fiber terminals in IML (left panels – arrows). NMDAR_GluN2B_ immunoreactivity is enriched in the IML of kainate–treated chronic animals (right panels – arrows) consistent with the localization of sprouted synapses in that subregion. Scale bars: 10 μm **b)** NMDAR–dependent sEPSCs are more abundant in GCs from TLE (bottom traces and graph) than from control mice (top traces and graph). Ifenprodil administration reduces the number of sEPSCs more in the GC from epileptiform (graph – light green) than from control DG (graph – dark green). **c)** Cumulative probability distribution of inter–event intervals from control (n = 150 consecutive events per each of 12 cells) (top graph) and epileptiform (n = 150 consecutive events per each of 13 cells) (bottom graph) GCs. In both cases, distributions of inter–event interval values were shifted rightward upon ifenprodil application. This is consistent with a decrease in sEPSC frequency in the presence of ifenprodil. However, the shift was grater in epileptiform GCs (p < 0.0001 for control neurons, p = 0.0001 for TLE GCs, KS–test) demonstrating the intensified contribution of ifenprodil– sensitive sEPSCs to the synaptic transmission in GC from TLE mice in comparison to neurons from control animals. **d)** Recordings of NMDAR–dependent sEPSCs from epileptiform granule cells revealed atypically large NMDAR–mediated synaptic currents characterised by amplitudes higher than 80 pA and lower than 500 pA (left trace – arrowheads) that were never observed in control GC (graphs). In rare cases, unusually large NMDAR–mediated EPSCs were observed in bursts (right trace).

**Fig 3.**
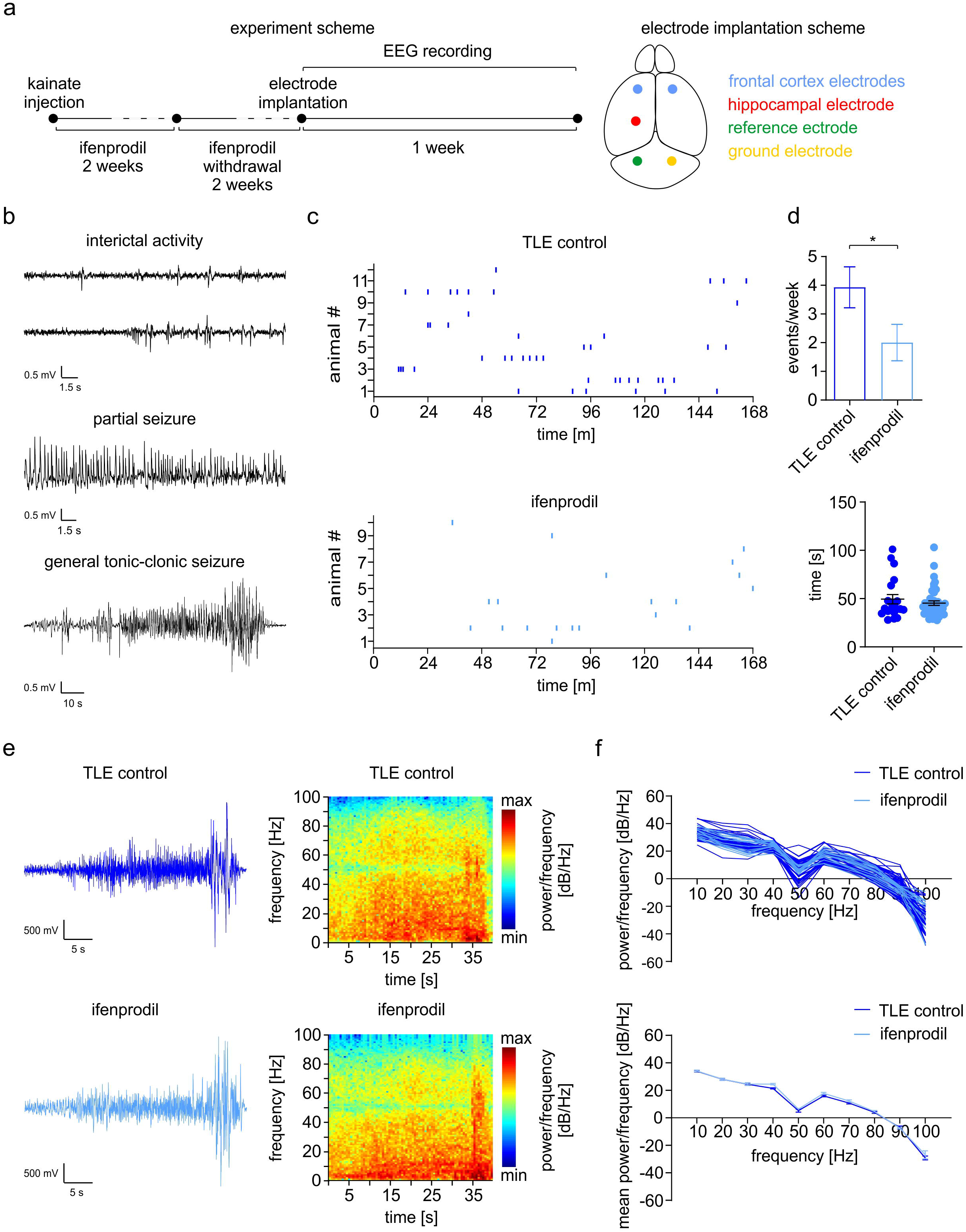
Ifenprodil administraton during epileptogenesis reduces the number of recurrent seizures. **a)** Plan of the experiment and the electrode implantation scheme. **b)** EEG recordings from freely moving TLE mice permits registration of distinct forms of synchronous epileptiform activity in the hippocampus including ictal tonic–clonic seizures (bottom trace). **c)** Distribution of recurrent seizure onsets over a week in control TLE animals (top graph) and TLE animals that received ifenprodil treatment during first two weeks of epileptogenesis (bottom graph). **d)** The average number of seizures per week was significantly reduced in animals that received ifenprodil treatment (n = 12) when compared to the control group (n = 10) (top graph – p = 0.046, MW–test). On the other hand, the seizure duration remained unchanged upon ifenprodil treatment (n = 43) in comparison to control conditions (n = 20) (bottom graph – p = 0.38, MW–test). **e)** Examples of electrographic tonic–clonic seizures, together with their spectrograms, recorded from control TLE animals (top panels) and ifenprodil–treated TLE animals (bottom panels). **f)** Ifenprodil treatment during the first two weeks of epileptogenesis does not alter magnitude of recurrent seizures as demonstrated by plots presenting spectra of power densities (top graph) and their means (bottom graph – p = 0.97, 2–way ANOVA).

**Fig 4.**
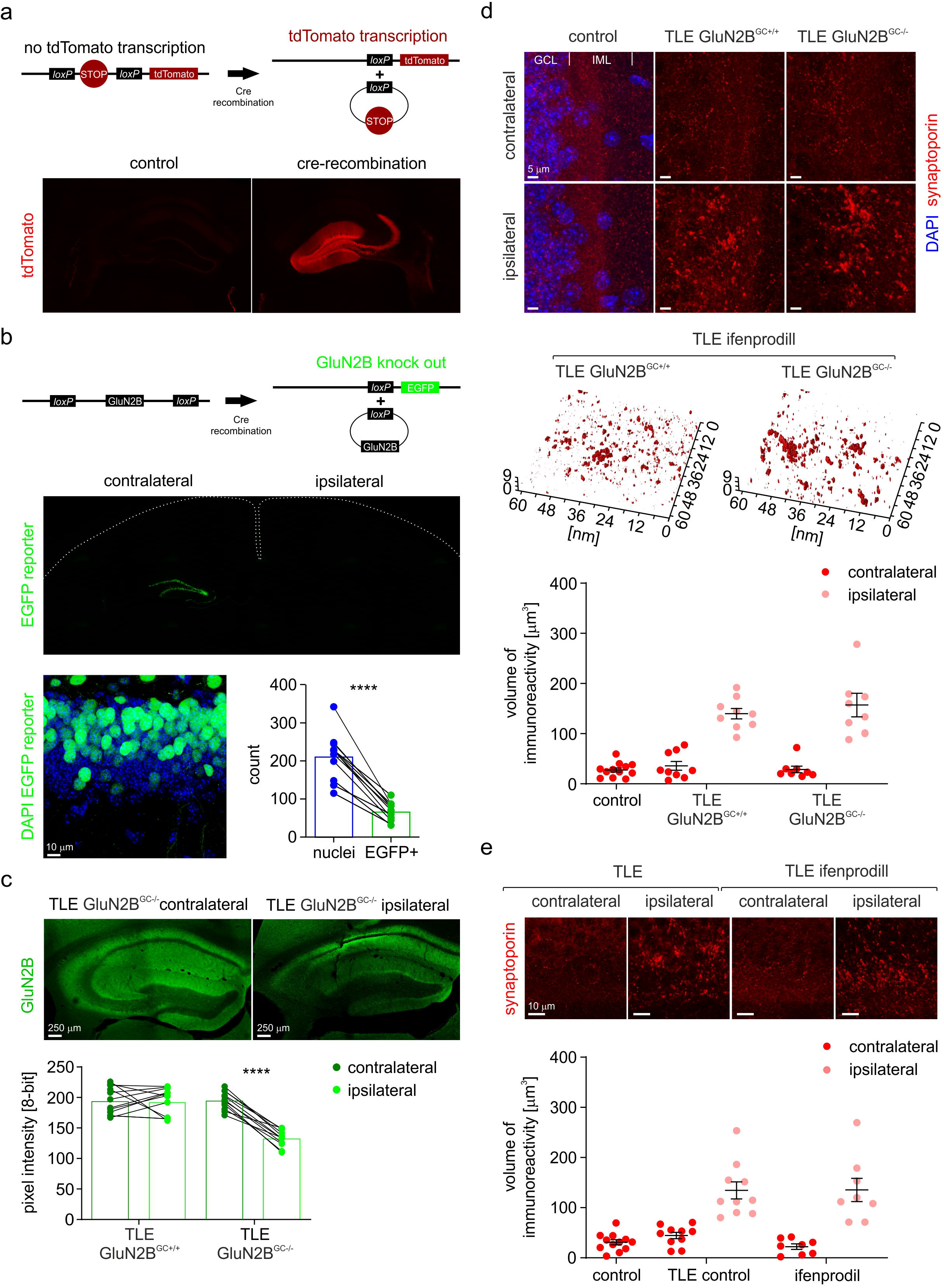
Mossy fiber sprouting in GluN2B–deficient animals. **a)** tdTomato expression was induced locally in the DG of transgenic mice (n = 3) in Cre–dependent fashion under control of C1ql2 promoter (scheme) to validate selectivity of viral gene transfer for dentate GC. Two weeks after the viral gene transfer, we observed a tdTomato–derived fluorescence that was strictly limited to the receptive field and projections of GCs at the injection site (ipsilateral) but not at the contralateral site (panels). Scale bars: 400 μm **b)** Efficacy of GC infection was estimated based on an eGFP reporter introduced in conjunction with GluN2B deletion (scheme) into the GC of GluN2B^ff^ mice (n = 12). Neuronal nuclei and eGFP– positive cells count in DG demonstrates that around 32 % of GCs were successfully infected with viral vector harboring the reporting gene (graph – p < 0.0001, t–test). Scale bars: top panel 500 μm, bottom panels 10 μm **c)** Brain sections from kainate–treated GluN2B^GC−/−^ animals (n = 12) and from kainate treated control GluN2B^GC+/+^ (GluN2B^f/f^) animals (n = 12) were incubated with anti GluN2B antibodies to map NMDAR_GluN2B_ expression four weeks after the epileptogenesis initiation by *status epilepticus*. Comparison of GluN2B staining intensities at the contralateral and ipsilateral site of GluN2B^GC−/−^ mice demonstrates that NMDAR_GluN2B_ expression in the molecular layer of DG was depleted as a result of successful gene transfer to the ipsilateral DG (panels and graph, p < 0.0001,t–test). GluN2B staining intensity in the molecular layer at the contralateral site of GluN2B^GC−/−^ animals did not differ significantly from that observed in control GluN2B^GC+/+^ mice (graph, p = 0.91, t–test). Scale bars: 250 μm **d)** Brain sections from kainate treated GluN2B^GC−/−^ animals (n = 8), kainate treated GluN2B^GC+/+^ animals (n = 9), and naive GluN2B^GC+/+^ (GluN2B^f/f^) animals (n = 12) were subjected to anti– synaptoporin immunodetection 4 weeks after the convulsant administration. Notable clusters of synaptoporin immunoreactivity were observed in the IML at ipsilateral sites of kainate–treated GluN2B^GC+/+^ and GluN2B^GC−/−^ mice but not in GluN2B^GC+/+^ specimens and contralateral sites of kainate treated GluN2B^GC−/−^ and GluN2B^GC+/+^ animals (top panels). There was no significant difference in the volume of immunoreactivity between the kainate–treated GluN2B^GC+/+^ and GluN2B^GC−/−^ mice (3D reconstructions and graph, p = 0.49, t–test). Scale bars: 5 μm **e)** Bran sections from naive (control), **s**aline–treated (TLE control group, n = 10) and ifenprodil– treated (ifenprodil group, n = 8) TLE animals one month after the intrahipocampal convuslsant administration were subjected to immunohistochemical synaptoporin detection. Notable clusters of synaptoporin immunoreactivity were observed only in the IML at ipsilateral sites of control TLE and ifenprodil–treated TLE mice, but not in control specimens and contralateral sites of control TLE and ifenprodil–treated TLE animals (panels). No significant difference in the volume of synaptoporin immunoreactivity in the IML was found between control TLE and ifenprodil–treated TLE mice (3D reconstructions and graph, p = 0.96, t–test). Scale bars: 10 μm

### Ifenprodil systemic administration

Animals were intraperitoneally injected with saline ifenprodil (Tocris / Biotechne) solution (20 mg per kg of body weight) and injections were repeated every 8 hours.

### Image and signal analysis

Image and signal edition and quantitative analysis was done using ImageJ (National Institutes of Health), Matlab (Mathworks), Prism (GraphPad Software Inc.), and Clampfit (Molecular Devices) software. In order to eliminate biases that the experimenter might introduce to the analysis, the names of all raw data files were encrypted using ‘Blind Analysis Tools’ (ImageJ extension) until analysis was completed.

ImageJ was used to calculate intensities of synaptoporin and GluN2B immunoreactivities. Multiplanar confocal images of equal volumes were maximum–projected, then regions of interest (ROIs) corresponding to the different hippocampal domains were selected and subjected to analysis with built–in ImageJ functions for the calculation of mean pixel value within the ROIs. Matched intensities of synaptoporin and GluN2B immunoreactivities in IML were processed with the Prism ‘correlation’ function to calculate the linear relationship between the two sets of data and to measure its normalized covariance (Pearson coefficient).

Spontaneous synaptic currents were detected and analyzed using Clampfit software based on a preselected template. All detected events were visually inspected and those that did not match general criteria expected for synaptic currents were rejected from the analysis. A cubic third–order polynomial was fitted to data in order to estimate the number of EPSCs beyond calculated time points. Cumulative probability distribution plots were built using Prism by pooling 150 consecutive events sampled per cell.

The analysis of synaptoporin immunoreactivity clusters in the IML of DG was performed using ImageJ as follows: equal–volum stacks of 8–bit confocal planes were binarized with the isodata algorithm (Ridler and Calvard, 1978). Dissociated single pixels were removed from the images with binary erode and dilate operations and images were segmented with a watershed algorithm. Finally, the number of non–zero (white) pixels were calculated to compute the total volume of acquired fluorescent signal. 3D surface reconstructions of the exemplary data were performed using the Matlab ‘isosurface’ function.

The number of neuronal nuclei was calculated in the following way. 24–bit RGB images were first constructed. Equally–volumed stacks of 8–bit confocal planes containing DAPI staining were then selected as the blue channel, while red and green channels remained empty. Nuclei were marked manually using a circular paintbrush marker with a pure green color (RGB: 0, 255, 0). When all the nuclei were marked, RGB images were split into separate red, green, and blue channels. The number of markers in the green channel was then calculated by the ImageJ function ‘Analyse particles’. We considered only nuclei that matched characteristic features of DAPI stained neuronal nuclei such as diameter (~10 μM), shape (circular), low density of chromatin with distinctive nuclear bodies (Fig. 5b – arrowheads). Nuclei that did not match these criteria were considered as non–neuronal or pyknotic (Fig. 5b – arrows) and excluded from the analysis. 3D surface reconstructions of the exemplary data were performed using the Matlab ‘isosurface’ function.

**Fig 5.**
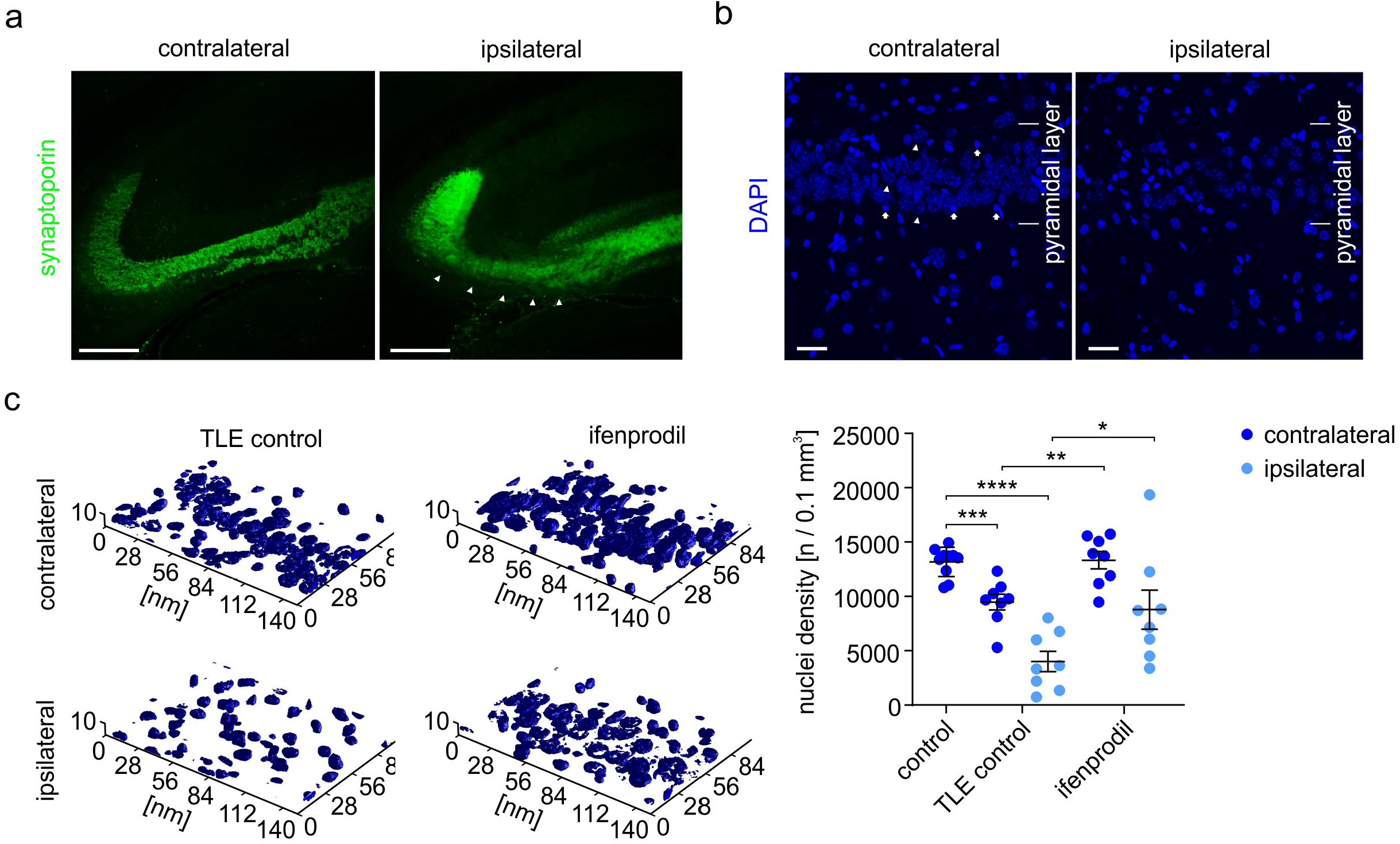
GluN2B block during epileptogenesis ensures neuroprotection. **a)** Confocal microscopy images of synaptoporin immunoreactivity in CA3 from TLE mice demonstrating neurodegenerative changes in mossy fiber pathway at the kainate injected (ipsilateral) hippocampus (right panel – arrowheads). Scale bars: 250 μm **b)** Confocal microscopy images demonstrating exemplary neurodegenerative changes in nuclei layout in a CA3 region of TLE mice. Neuronal nuclei were selected for quantification based on their characteristic features including circular shape, large size and presence of nuclear bodies (arrowheads), that clearly discriminate them from small, oval, with dense chromatin non–neuronal or pycnotic nuclei (arrows). Scale bars: 20 μm **c)** Quantification of neuronal density in the CA3 pyramidal layer of control TLE (n = 8) and ifenprodil–treated (n = 8) TLE animals demonstrates cell loss at both contralateral (graph – dark blue, p = 0. 0003, t–test) and ipsilateral (graph – light blue, p < 0.0001, t–test) sites in comparison to naive control (n = 10) specimens (graph – dark blue). Ifenprodil treatment during epileptogenesis limited neurodegenerative processes at contralateral (top panels and graph – dark blue, p = 0.003, t–test) and ipsilateral (bottom panels and graph – light blue, p = 0.033, t–test) sites of TLE animals.

The occurrence of seizures was evaluated off–line by visual inspection of the EEG and video recordings with Twin Software (Grass Technologies) by trained observers. An electroencephalographic seizure was defined as low–amplitude rhythmic activity discharges with polyspikes that lasted > 10 s and was associated with behavioral correlates visible on video monitoring. The beginning and the end of the electrographic seizure were defined as points between which the discharge amplitude exceeded at least twice the magnitude of background activity. For wavelet analysis, electrographic seizures were exported to Matlab (Mathworks) and processed with spectrogram(x) and fft(x) functions to compute spectrograms and calculate nonparametric power spectral density using fast Fourier transforms.

### Statistics and charts

Prism (GraphPad Software Inc.) and Matlab (Mathworks) were used for numeric data management and statistics. Prism was used to build graphs. In order to examine whether the data represent a normal distribution, we applied the one–sample Kolmogorov–Smirnov test. Either the parametric t–test or nonparametric Mann–Whitney U test (MW–test) were performed to investigate the null hypothesis that mean values come from the two data sets with the same distribution. Cumulative probability distributions were compared using the two–sample Kolmogorov–Smirnov test (KS–test). Mean spectra of power densities were compared using 2–way ANOVA for balanced designs.

Values in charts are given as means ± S.E.M. The result was considered statistically significant when p was < 0.05. Asterisk notation was used in order to flag the levels of significance (* p < 0.05; ** p < 0.01; *** p < 0.001; **** p < 0.0001). Figures were prepared with the Corel Package (Corel Corporation).

## Results

### Hippocampal GluN2B expression in TLE is upregulated in recurrent mossy fiber terminals

In the hippocampus, axons of dentate granule cells (GCs) –mossy fibers– create functional connections with CA3 pyramidal neurons via presynaptic terminals known as giant mossy fiber boutons. These giant mossy fiber boutons form a specific morphological pattern, histologically distinguished as *stratum lucidum*. In an attempt to elucidate the role of GluN2B in the pathophysiology of TLE, we first examined NMDAR_GluN2B_ expression in the mossy fiber pathway in hippocampal brain sections from control (saline treated) and TLE mice treated with pilocarpine (a muscarinic agonist commonly used to promote epileptogenesis via the precipitating seizure insult called *status epilepticus*) (Turski et al., 1984). We immunodetected the GluN2B subunit (selectivity of the antibody was validated elsewhere) (Fukushima et al., 2009) and synaptoporin, a marker of giant mossy fiber boutons, which is a splice variant of synaptophysin predominantly expressed in these synaptic terminals.

The altered NMDAR_GluN2B_ immunoreactivity was observed in multiple hippocampal domains eight weeks after the pilocarpine treatment (Fig. 1a, top panels). In particular, we detected increased GluN2B immunoreactivity in the inner molecular layer (IML) of DG (Fig. 1a, arrows in top right panel). In contrast, in control conditions NMDAR_GluN2B_ expression was uniformly distributed over the receptive field of dentate GCs (Fig. 1a, left panel). To quantify this observation, we calculated the mean fluorescence intensities of GluN2B immunoreactivity in different DG subregions in brain sections from control (n = 8) and TLE animals (n = 8) eight weeks after pilocarpine administration (Fig. 1a, graph). We observed significant upregulation of NMDAR_GluN2B_ in the epileptiform GC layer (GCL) (mean pixel value 26.9 ± 2.54 for control and 51.25 ± 2.41 for TLE animals, p < 0.0001, t–test) and IML (mean pixel value 87.73 ± 4.77 for control and 132.50 ± 3.98 for TLE animals, p < 0.0001, t– test). On the other hand, NMDAR_GluN2B_ was downregulated in the epileptiform hilus (mean pixel value 34 ± 0.75 for control and 23.77 ± 0.58 for TLE animals, p < 0.0001, t–test) and outer molecular layer (OML) (mean pixel value 82.61 ± 4.52 for control and 64.79 ± 4.36 for TLE animals, p = 0.013, t– test). This abnormal NMDAR_GluN2B_ expression pattern was observed as early as four weeks after the induction of *status epilepticus* and was still maintained twelve weeks after the insult (Fig. 1a, bottom panels).

The observed NMDAR_GluN2B_ expression in epileptiform DG is spatially restricted with very distinguishable laminar distribution in the IML. This is typically attributed to sprouted mossy fiber terminals. Sprouting of hippocampal mossy fibers is one of the best–described, seizure–evoked events which takes place during epileptogenesis in both human patients and animal models of TLE (Scharfman et al., 2003). Sprouted mossy fibers create feedback connections on the GCs from which they originate, forming aberrant synapses in the IML and, less prominently, GCL. We noted an extensive sprouting of mossy fibers in the DG of TLE mice (n = 8) eight weeks after the convulsant administration, as indicated by synaptoporin immunoreactivty in the IML of DG (Fig. 1b, bottom panels, arrows in bottom left panel). Such immunoreactivity was imperceptible in control conditions (n = 8) (Fig. 1b, top panels). Morphometric analysis of the mean immunostaining intensity across the DG (Fig. 1b, graph) confirmed this observation: a significant upregulation of synaptoporin expression in the epileptiform GCL (mean pixel value 23.06 ± 1.48 for control and 64.76 ± 4.36 for TLE animals, p < 0.0001, t–test) and IML (mean pixel value 34.98 ± 1.70 for control and 93.65 ± 8.68 for TLE animals, p < 0.0001, t–test). Synaptoporin expression remained unaltered in the epileptiform hilus (mean pixel value 137.8 ± 4.62 for control and 145.5 ± 3.62 for TLE animals, p = 0.21, t–test) and OML (mean pixel value 15.28 ± 1.02 for control and 18.38 ± 1.81 for TLE animals, p = 0.15, t–test).

The spatial pattern of increased synaptoporin immunoreactivity in the IML of pilocarpine– treated mice thus corresponds to the localization of increased NMDAR_GluN2B_ immunoreactivityin the same region and model (compare Fig. 1a, and Fig. 1b – arrows; compare graphs in Fig. 1a and Fig. 1b). Moreover, a correlation in time between the rise of these two signals was clearly visible in tested brain slices. Sprouting intensified as TLE developed (Fig. 1c, panels) and this was associated with a gradual increase of NMDAR_GluN2B_ expression in the IML during epileptogenesis (Fig. 1a, bottom panels) as demonstrated by the linear correlation between the synaptoporin and the NMDAR_GluN2B_ expression at different time points (n = 4 × 8) (Fig. 1c, graph) (Pearson correlation coefficient r = 0.74). Collectively, our results demonstrate that in the pilocarpine–based model of TLE, expression of a specialized subpopulation of NMDAR_GluN2B_ is altered in the hippocampus and corresponds to the mossy fiber feedback connectivity at GCs.

### GluN2B–mediated excitatory drive onto DG granule cells is altered in TLE

As GluN2B may play an important role in pathophysiology of the epileptiform DG, we decided to investigate the functional impact of this subunit on GC physiology in control conditions and in TLE. We used the TLE model based on unilateral intrahippocampal kainate injection as it most reliably mimics different aspects of human pathology (Bouilleret et al., 1999). Kainate–treated animals develop chronic convulsions within a few weeks upon convulsant–induced seizure insult. Chronic changes at the injection site involve neuroinflammation, cell death, dispersion of dentate GCs, and far–reaching alterations in synaptic connections and their receptor composition, including mossy fiber sprouting. The change of model also enabled us to investigate whether epileptiform NMDAR_GluN2B_ alterations are universal in different models of TLE.

First, we checked whether the NMDAR_GluN2B_ expression pattern corresponds to mossy fiber sprouting in the DG of kainate–treated animals. For this purpose, immunohistochemical detection of GluN2B and synaptoporin four weeks after the convulsant injection into the left hippocampus and the consequent *status epilepticus*, was carried out. As in the pilocarpine model, an extensive sprouting of mossy fibers was observed in the DG of kainate–treated mice as indicated by the presence of synaptoporin immunoreactivity clusters in the IML of ipsilateral site (Fig. 2a, left panels, arrows). In accordance with synaptoporin immunoreactivity, we also observed increased GluN2B expression with a laminar distribution in the IML, corresponding to the localization of synaptoporin–positive sprouted synaptic terminals (Fig. 2a, right panels, arrows).

Next, we compared the extent of NMDAR activation by the quantal release of glutamate at the control and epileptiform dentate GCs. For that purpose, we recorded TTX–sensitive, spontaneous excitatory post–synaptic currents (sEPSCs) from the GCs in hippocampal brain sections from control and TLE animals (ipsilateral hemisphere). NMDAR–dependent sEPSCs were evoked by a +40 mV holding potential and isolated through the continuous presence of bicuculline (10 mM), CGP (3 μM), and NBQX (40 mM) in extracellular medium. Under these conditions, we observed sEPSCs which were mediated by NMDAR receptors, because they were completely abolished with the NMDAR antagonist D–AP5 (data not shown). However, the frequency of those NMDAR–mediated sEPSCs was insufficient for conclusive analysis. Therefore, we decided to increase the spontaneous synaptic activity in the recorded cells using 4–AP (50 μM), a potassium channel antagonist. In the presence of 4–AP (added immediately after the beginning of the recording) the frequency of NMDAR–mediated sEPSCs in control (n = 12) and TLE (n = 13) GCs gradually increased and, 10 minutes after the drug infusion, was significantly lower in control GCs than in GCs from TLE mice (Fig. 2b, left traces, ‘reference activity’ in the graph).

In order to determine the contribution of the NMDAR_GluN2B_ subtype to the total NMDAR– dependent conductance, we applied ifenprodil (5 μM) 15 minutes after the beginning of the recording and 4–AP administration (Fig. 2b – graph). We noticed that upon saturation with ifenprodil (10 minutes after the antagonist application) the number of NMDAR_GluN2B_–dependent sEPSCs per minute was slightly reduced in control neurons, while in TLE specimens the reduction was much more prominent (Fig. 2b, right traces, ‘ifenprodil’ in the graph). This is further illustrated by comparison of cumulative probability distributions of inter–event intervals during 5–minute reference activity (n=150 consecutive events sampled per cell) and upon saturation with ifenprodil (n = 150 consecutive events sampled per cell) in both control (Fig. 2c – top graph) and TLE (Fig. 2c – bottom graph) GCs. After infusion of ifenprodil to the extracellular solution, cumulative probability distributions of sEPSC inter–event intervals from both investigated groups exhibited a rightward shift relative to reference activity indicating a decrease in the frequency of NMDAR–conducted sEPSCs (p < 0.0001 for control GCs and p < 0.0001 for TLE GCs, KS–test). However, the shift was larger in epileptiform GCs, demonstrating greater contribution of ifenprodil–sensitive sEPSCs to the synaptic transmission in the GC in TLE mice compared to neurons in control animals.

Interestingly, in some cases (5 out of 13 cells) we observed unusually large (Fig. 2d, left trace, arrowheads) NMDAR–mediated sEPSCs characterized by an amplitude which considerably exceeded the amplitude of typical (Fig. 2d – left trace, asterisks) NMDAR–mediated sEPSCs. Those large postsynaptic currents were never observed in control conditions. In very rare cases (2 out of 5 cells), they were released in bursts (Fig. 2d – right trace). The distributions of recorded NMDAR–mediated sEPSC amplitudes from the control and TLE GCs clearly demonstrate a separate population of currents, characterized by magnitudes over 80 pA, which were conducted (during ‘reference activity’) in TLE GCs but not in control neurons (Fig. 2d – graphs). These large events were also observed under influence of ifenprodil (data not shown). Although the phenomenon was not further investigated, our observations emphasize the special role of the NMDAR receptor in the pathophysiology of epileptiform DG.

### Prolonged ifenprodil administration during epileptogenesis reduces the number of chronic seizures in mice with TLE

Seizures are abnormal hypersynchronous discharges (lasting tens of seconds) of a group of neurons. In TLE, seizures are recurrent and they originate from limbic structures such as the hippocampus. The electrographic signature of seizures are evident in EEG recordings and resemble the major clinical symptom of TLE (Bertram, 2009; Jefferys et al., 2012; Jiruska et al., 2013). Knowing that NMDAR_GluN2B_ plays an important role in pathophysiology of hippocampal networks in TLE, we investigated whether selective GluN2B antagonism might interfere with epileptogenesis using vEEG recordings.

We subjected kainate–treated mice to either intraperitoneal ifenprodil (experimental group, 20 mg per kg of body mass) or saline (control group) injections for two weeks in eight–hour intervals. The treatment started the next day after the convulsant administration and was terminated after two weeks. Two weeks later the electrodes were implanted into the hippocampi and the animals were subjected to one–week vEEG recording (drug–free) (Fig. 3a – experiment and electrode implantation scheme).

In all saline and ifenprodil treated animals, we observed periods of slow–wave interictal activity with isolated spikes and spike clusters (Fig. 3b – top pair of traces) which typically occur in intrahippocampal EEG recordings in mice (Twele et al., 2017). In addition, in most of the recorded animals, we noticed paroxysmal activity characterized by high–amplitude trains of spikes (Fig. 3b – middle trace). These trains of spikes resemble a partial non–convulsive seizure (Twele et al., 2017). Moreover, in ~60 % of tested mice, we recorded ictal electrographic seizures resembling a generalized convulsive seizure (Twele et al., 2017), which started as low–amplitude and high–frequency activity that gradually evolved into multiple spikes (Fig. 3, bottom trace). On simultaneous video recordings, ictal electrographic seizures were manifested as tonic–clonic convulsions with loss of posture preceded and followed by short periods of immobility. Only recordings from animals that experienced at least one generalized convulsion confirmed by both EEG and video were subjected to the analysis. We determined the seizure onset profile for each animal (Fig. 3c) and observed a reduction in the number of seizures in ifenprodil–treated TLE mice (n = 12) compared to control TLE mice (n = 10). The seizures occurred at an average rate of 3.91 ± 0.71 per week in control TLE animals and 2.00 ± 0.63 in ifenprodil–treated TLE animals (p = 0.046, MW–test) (Fig. 3d – top graph) without any significant difference in their duration between the groups (n = 43 events for control TLE animals and n = 20 events for ifenprodil–treated TLE mice, p = 0.38, t–test) (Fig. 3d – bottom graph). Wavelet analysis showed that the average power of the analyzed seizures did not vary between groups, either as demonstrated by exemplary traces and their spectrograms (Fig. 3e), or by spectral power densities (Fig. 3f – top graph) and their means (p = 0.97, 2–way ANOVA) (Fig. 3f – bottom graph). Taken together, the results demonstrate that prolonged ifenprodil–mediated antagonism of NMDAR_GluN2B_ during epileptogenesis downregulates the number of seizures by ~46 % during the chronic phase of TLE, without any significant effect on seizure magnitude or duration.

### NMDAR_GluN2B_ inhibition during epileptogenesis does not prevent formation of mossy fiber sprouting

The role of GCs’ recurrent connectivity in epileptogenesis is controversial. Mossy fiber sprouting constitutes both an aberrant positive–feedback at GCs (Scharfman HE, Sollas AL, Berger RE; Wuarin and Dudek, 1996; Buckmaster et al., 2002), which is suggested to be seizurogenic (Feng et al., 2003; Tauck and Nadler, 1985), and excitatory synaptic input to inhibitory interneurons in DG, which possibly exerts anti seizurogenic effect (Sloviter, 1992; Sloviter et al., 2006). Despite the ambiguous role that sprouting plays in the epileptiform hippocampus, more and more data confirm that this phenomenon greatly contributes to the onset of chronic seizures, stimulating granule cells when excitation counterbalances inhibition in the DG network (Sutula and Dudek, 2007). We have already identified NMDAR_GluN2B_ as a possible component of sprouted mossy fiber synapses and demonstrated that GluN2B–aimed antagonism during epileptogenesis is sufficient to diminish the number of chronic seizures. As synaptic activity in developing connections promotes the formation of synapses (Flavell and Greenberg, 2008), we investigated whether NMDAR_GluN2B_ antagonism blocks mossy fiber sprouting during epilepsy development. We took advantage of cell–specific gene deletion using C1ql2–driven expression, which is largely restricted to the dentate GCs in the hippocampus (Iijima et al., 2010). The C1ql2 promoter was used in a lentiviral construct to provide the expression of Cre recombinase and eGFP reporter selectively to GCs. The validity of our approach was first tested in mice with Cre–dependent tdTomato expression (n = 3) (Fig. 4a, scheme). Two weeks after the viral gene transfer through intrahippocampal vector administration, we acquired a tdTomato– derived fluorescence strictly limited to the receptive field and projections of GCs. (Fig. 4a, images). Then, we deleted GluN2B in GCs during epileptogenesis using GluN2B^ff^ mice (GluN2B^GC−/−^, n = 12). The animals were first subjected to unilateral (left hemisphere) intrahipocampal vector injection and two weeks later to the intrahippocampal kainate administration to the same hemisphere. Four weeks after the convulsant treatment, we estimated the efficacy of infection comparing the number of eGFP expressing neurons in the GC layer of GluN2B^GC−/−^ mice (Fig. 4b – scheme) to the total number of neuronal nuclei in exactly the same region and tissue volume. We found that on average 31.8 ± 2.5 % of GCs were successfully infected with the viral vector (the mean number of neuronal nuclei was 213.6 ± 17.36 and the mean number of GFP–positive cells was 67.67 ± 6.931, p < 0.0001, t–test) (Fig. 4b –graph).

Further, brain sections from the same kainate–treated GluN2B^GC−/−^ animals (n = 12) and from appropriate control (GluN2B^GC+/+^) mice (kainate–treated GluN2B^f/f^ animals) (n = 12) were incubated with anti GluN2B antibodies and the subunit immunoreactivity was analyzed in both groups to determine whether NMDAR_GluN2B_ expression was affected by introduction of the construct into the GCs, four weeks after the epileptogenesis initiation by *status epilepticus*. When comparing GluN2B staining intensities at the contralateral and ipsilateral site of TLE GluN2B^GC+/+^ and TLE GluN2B^GC−/−^ mice, we noticed that NMDAR_GluN2B_ expression in the ipsilateral DG molecular layer of GluN2B^GC−/−^ animals was, on average, depleted by 31.57 ± 2.22 % (the mean 8–bit pixel value was reduced from 195.3 ± 4.15 at collateral site to 133.2 ± 3.86 at ipsilateral site, p < 0.0001, t–test) (Fig. 4c panels and graph). The GluN2B staining intensity in the DG molecular layer at the contralateral site of GluN2B^GC−/−^ animals did not differ significantly from that observed in the same region at the contralateral and ipsilateral sites of TLE GluN2B^GC+/+^ mice (the mean 8–bit pixel value equaled 195.3 ± 4.15 at GluN2B^GC−/−^ contralateral site and 194.4 ± 6.48 at GluN2B^GC+/+^ contralateral site, p = 0.91, t– test; the mean 8–bit pixel value equaled 193.4 ± 6.45 at GluN2B^GC+/+^ ipsilateral site, p = 0.81, t–test) (Fig. 4c – graph). Finally, brain sections from TLE GluN2B^GC−/−^ animals (n = 8), TLE GluN2B^GC+/+^ animals (n = 9) were subjected to anti–synaptoporin immunodetection 4 weeks after convulsant administration to measure the influence of GluN2B downregulation on sprouting formation during epileptogenesis and confront that results with data obtained alike from naive GluN2B^GC+/+^ (GluN2B^f/f^) animals (n = 12). The extent of mossy fiber sprouting in the IML was measured in brain sections as a volume covered by synaptoporin immunoreactivity. In contrast to the GluN2B^GC+/+^ specimens and contralateral sites of kainate treated GluN2B^GC−/−^ and GluN2B^GC+/+^ animals, the clusters of synaptoporin immunoreactivity were observed in the IML at ipsilateral sites of kainate–treated GluN2B^GC+/+^ and GluN2B^GC−/−^ mice (Fig. 4d – top panels) without any significant difference in the volume of immunoreactivity between the two latter groups (139.9 ± 10.28 μm^3^ for kainate treated GluN2B^GC+/+^ and 157.0 ± 23.40 μm^3^ for kainate treated GluN2B^GC−/−^, p = 0.49, t–test) (Fig. 3d – 3D reconstructions and graph).

Subsequently, we applied the same experimental protocol to genetically unaltered (C57BL6) ifenprodil–treated (ifenprodil group, n = 8), saline–treated (TLE control group, n = 10) TLE animals, one month after the intrahippocampal convulsant administration (animals received either saline or ifenprodil intraperitoneal injections for two weeks following kainate treatment as in the previous experiments). The same experimental protocol was also applied to untreated (C57BL6) (control group, n = 12) mice. As in our previous experiments, the data show the clusters of synaptoporin immunoreactivity only in IML at ipsilateral sites of control TLE and ifenprodil–treated TLE mice but not in control specimens and contralateral sites of control TLE and ifenprodil–treated TLE animals (Fig. 4e, panels). No significant difference in the volume of synaptoporin immunoreactivity in IML was found between control TLE and ifenprodil–treated TLE mice (134.4 ± 17.01 μm^3^ for control TLE and 135.4 ± 23.13 μm^3^ for ifenprodil–treated TLE, p = 0.96, t–test) (Fig. 4e, 3D reconstructions and graph). Together, our findings demonstrate that neither cell–specific partial deletion of NMDAR_GluN2B_ in the hippocampus nor systemic pharmacological inhibition of NMDAR_GluN2B_ function does affect mossy fiber sprouting formation.

### NMDAR_GluN2B_ inhibition during epileptogenesis reduces deleterious neurodegenerative changes

Seizure leads to excitotoxicity and activation of self–destructing neurodegenerative pathways as a result of, i.a., persistent depolarization of neurons followed by mitochondrial and energy failure, increased production of reactive oxygen species, and increased concentration of intracellular Ca^2+^ (Favaron et al., 1990; Mills and Kater, 1990). Consequently, neurodegenerative changes are frequently observed in patients with acquired epilepsy (Mathern et al., 1995) and in different animal models of TLE, including kainate–induced epileptogenesis (Wang et al., 2008). While it is clear that seizures result in excitotoxic neurodegeneration, it is not clear whether or not neurodegenerative changes are potentially pro–epileptogenic. Nevertheless, some experimental approaches targeting neurodegenerative pathways have been effective against epileptogenesis (Jones et al., 2012; Van Vliet et al., 2012; Liu et al., 2016; Semple et al., 2017).

The glutamatergic activation of postsynaptic NMDARs leads to increased intracellular Ca^2+^ concentrations, which suggests that NMDAR_GluN2B_ may be involved in excitotoxic neuronal death (Bullock et al., 1992; Zhou and Baudry, 2006). To test this hypothesis, we took advantage of the high vulnerability of hippocampal CA3 neurons to degeneration in the kainate model of epilepsy, and inspected whether NMDAR_GluN2B_ antagonism exerts a neuroprotective effect in our experimental approach.

We counted neuronal nuclei in a pyramidal layer to compare the sensitivity of the CA3 hippocampal region to neurodegeneration in brain sections from saline–treated (TLE control group, n = 8) and ifenprodil–treated (ifenprodil group, n = 8) TLE animals one month after intrahippocampal convulsant administration (animals received intraperitoneal injections of either saline or ifenprodil for two weeks following kainate treatment as in the previous experiments). We compared the results obtained in TLE mice with samples from the naive controls (control group, n = 10). Prior to the analysis, the hippocampal brain sections were subjected to the fluorescent synaptoporin immunodetection and co–stained with DAPI.

In TLE animals, we commonly observed maformed synaptoporin immunoreactivity (Fig. 5a – arrowheads) in the CA3 at the site of kainate injection which indicates CA3 sclerosis. Only brains showing signs of that neurodegenerative changes were further examined. Neuronal nuclei were identified based on their discriminative features revealed by DAPI staining (Fig. 5b, arrowheads), which allowed for a reliable exclusion of non–neuronal nuclei (Fig. 5b, arrows). We confirmed neurodegeneration in the CA3 region of convulsant–treated mice at both the contralateral (9485 ± 734 nuclei per 0.1 mm^3^ for TLE animals in comparison to 13189 ± 428 nuclei per 0.1 mm^3^ for control subjects, p = 0. 0003) and ipsilateral site (4007 ± 940 nuclei per 0.1 mm^3^ for TLE animals in comparison to 13189 ± 428 nuclei per 0.1 mm^3^ for control subjects, p < 0.0001, t–test) (Fig. 5c – graph). Importantly, administration of ifenprodil during epilepsy development markedly reduced the severity of cell death in the CA3 region at both contralateral (13347 ± 802 nuclei per 0.1 mm^3^ for ifenprodil–treated TLE animals and 9485 ± 734 nuclei per 0.1 mm^3^ for control TLE animals, p = 0.003, t–test) and ipsilateral site (13189 ± 428 nuclei per 0.1 mm^3^ for ifenprodil–treated TLE animals and 8798 ± 1802 nuclei per 0.1 mm^3^ for control TLE animals, p = 0.033, t–test) (Fig. 5c –panels and graph). Hence, we conclude that GluN2B–containing NMDARs contribute to seizure induced neuronal cell death in epileptogenesis.

### Ifenprodil treatment inhibits transition of epileptiform hippocampal network into seizure

Next, we examined whether the NMDAR_GluN2B_ antagonism might be effectively used to target ictogenesis. We first investigated the influence of ifenprodil on the evolution of network synchronization during conversion from interictal activity into seizure *ex vivo* (Dossi et al., 2014; Gong et al., 2014; Chang et al., 2019) by inspecting progressive changes in local field potential in the DG. The epileptiform discharges were induced in hippocampal slices obtained from TLE mice one month after kainate treatment. Slices were randomly divided into two groups: control and ifenprodil– treated (the sections were continuously superfused with extracellular solutions containing 5 μM ifenprodil). Each slice was visually inspected with a microscope prior to the experiment to ensure slice integrity and the presence of CA3 sclerotic atrophy – one of the morphological hallmarks of epileptic network disorganization. Finally, sections were transferred to the interface chamber and superfused for 40 minutes with K^+^–enriched artificial cerebrospinal fluid (ACSF_>K_) which was then exchanged with K^+^– enriched and Mg^2+^–free artificial cerebrospinal fluid (ACSF_>KØMg_) for another 40 min (Fig. 6a – experiment scheme). Both extracellular solutions also contained bicuculline (10 μM) and CGP (3 μM) in order to pharmacologically block inhibitory synaptic transmission.

**Fig 6.**
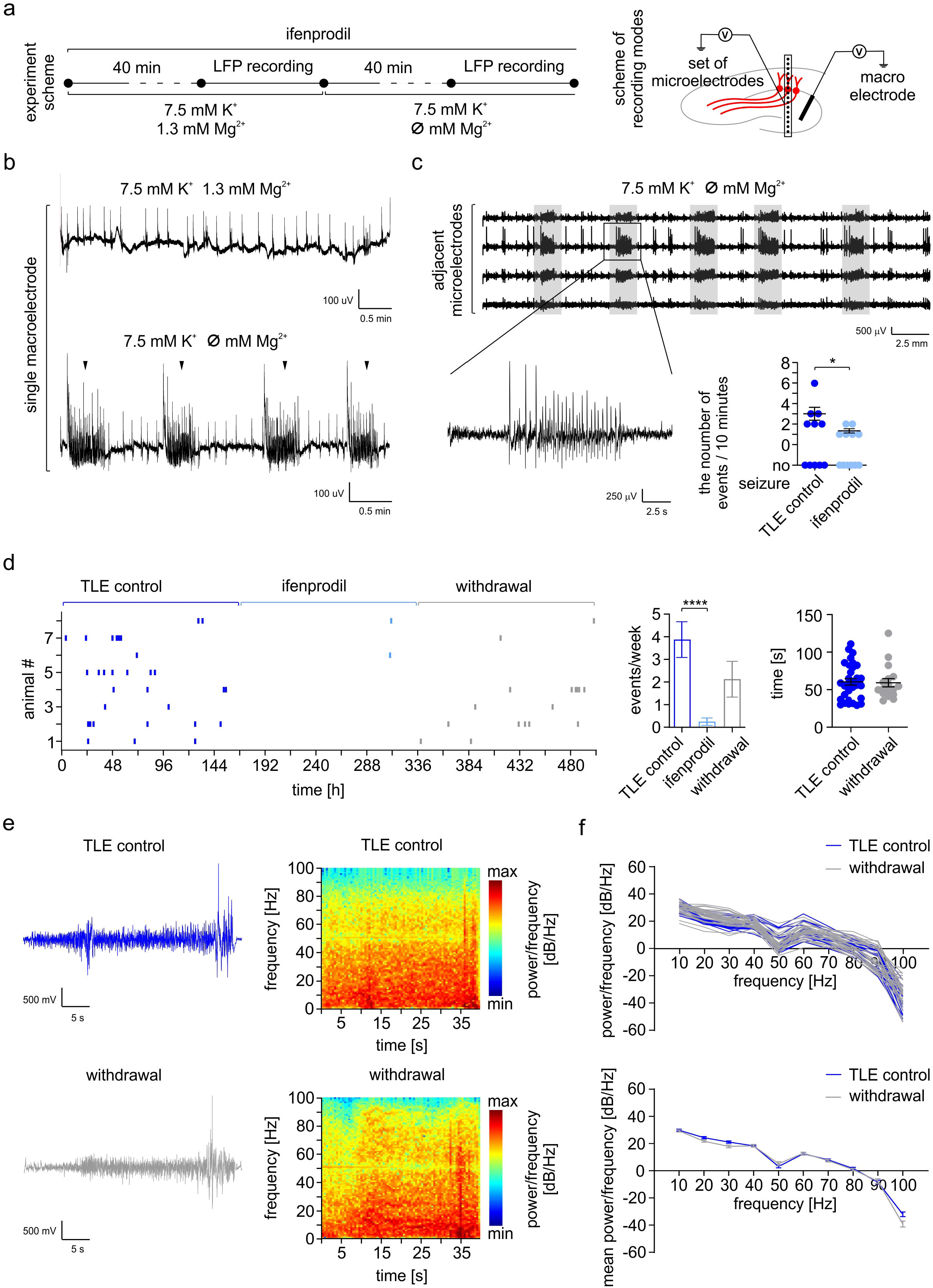
Ifenprodil administration efficiently blocks ictogenesis *ex vivo* and *in vivo*. **a)** Plan of the experiment and schemes of the recording modes. **b)** Exemplary recordings from the DG of TLE brain slices perfused with ACSF_>K_ show spontaneous interictal–like activity, exhibiting biphasic local field potential shifts lasting several tens of milliseconds (top trace). Exchange of perfusion medium with ACSF_>KØMg_ provokes ictal–like events in the DG of TLE brain slices (bottom trace – arrowheads). **c)** Comparison of the number of induced ictal–like events (zoomed portion of a trace) per 10 minutes in control (n = 6 out of 11) and ifenprodil–treated (n = 6 out of 12) TLE brain sections demonstrates that NMDAR_GluN2B_ inhibition reduced the probability of network transition from interictal to ictal activity (graph – p = 0.021, MW–test). **d)** Distribution of recurrent seizure onsets in TLE animals (n = 8) over three weeks of EEG recording that demonstrates considerable suppression of recurrent seizures during second week when ifenprodil was administrated (light blue). The average number of seizures per week was significantly reduced by ifenprodil administration in comparison to control conditions (left graph, p < 0.0001, MW–test). Ifenprodil withdrawal restored the seizure occurrence and the average number of seizures per week after withdrawal was not significantly different than during control conditions (left graph, p = 0.13, MW–test). Seizure duration remained unchanged upon ifenprodil withdrawal (n = 17) in comparison to control conditions (n = 31) (right graph, p = 0.86, t–test). **e)** Examples of electrographic seizures, together with their spectrograms, recorded from TLE animals under control conditions (top panels) and after ifenprodil withdrawal (bottom panels). **f)** Magnitude of seizures upon ifenprodil withdrawal (n = 17) was not significantly different from the magnitude of seizures recorded during control conditions (n = 31) as demonstrated by plots presenting spectra of power densities (top graph) and their means (bottom graph, p = 0.99. 2–way ANOVA).

While perfusing the TLE brain slices with ACSF_>K_ we acquired (using either single macroelectrode or set of microelectrodes – see scheme of recording modes in Fig. 6a) spontaneous interictal activity in the DG, in the form of biphasic local field potential shifts lasting several tens of milliseconds. A representative example of this activity is illustrated in the macroelectrode–recorded trace (Fig. 6b – top trace). The increase of K^+^ concentration to 7.5 mM was not enough to induce seizures in brain sections even when the network was disinhibited with bicuculline and CGP. Nevertheless, after 40–minute perfusion with ACSF_>KØMg_ we recorded the ictal–like events in the DG of TLE brain slices, as can be seen in the macroelectrode recording (Fig. 6b – bottom trace, arrowheads) and adjacent microelectrode recordings (Fig. 6c). The evoked ictal–like activity was usually composed of biphasic local field potential deflections with embedded multi–unit spikes (Fig. 6b, c). The successful transition from the interictal into ictal state was observed in 50 % of control and 58.3 % of ifenprodil–treated samples (Fig. 6c –left graph). Importantly, when AP–5 was present in the extracellular solution, ictal–like events were never observed (data not shown), indicating that conversion from a seizure prone hippocampal network into seizure is NMDAR–dependent in our protocol. When we compared the number of ictal–like events per 10 minutes of recording in control TLE (n = 6) and ifenprodil treated (n = 6) TLE sections, we found a reduction of event frequency upon NMDAR_GluN2B_ inhibition (from 3 ± 0.63 for control conditions to 1.33 ± 0.21 when ifenprodil was present in the extracellular solution, p = 0.021, MW–test) (Fig. 6c – right graph). Taken together, the protocol we used allowed for generation of stable ictal events in the brain section which reliably reproduce the electrographic seizures recorded *in vivo*. Our data demonstrate that ifenprodil in an extracellular solution reduces the probability of seizure occurrence by ~66 % during ex vivo ictogenesis.

### Ifenprodil administration ameliorates seizure expression in mice with TLE

Having shown that NMDAR_GluN2B_ antagonism modulates ictogenesis via reduction of seizure probability in the epileptiform hippocampal network *ex vivo*, we next investigated whether GluN2B antagonism might also suppress recurrent seizures in TLE *in vivo*. Mice that had been treated with kainate were subjected to one–week vEEG recordings one month after convulsant administration. Ictal electrographic seizures resembling a generalized convulsive seizure occurred in ~ 60% of recorded animals. Animals that had experienced at least one generalized convulsion confirmed with both EEG and video recording (n = 8) were then subjected to intraperitoneal ifenprodil administration (20 mg per kg of body mass) for one week in eight–hour intervals. After that, ifenprodil was withdrawn and the recordings were continued for one more week.

We then determined the seizure onset profile for each treated animal (Fig. 6d, left graph) and noted that 6 out of 8 mice with initial chronic seizure (dark blue) were completely seizure–free during the entire ifenprodil administration period (light blue). However, in all these animals, recurrent epileptic discharges reappeared immediately upon ifenprodil withdrawal (grey). The average occurrence of seizure was reduced from 3.87 ± 0.78 per week in control TLE to 0.25 ± 0.16 per week when ifenprodil was administered (Fig. 6d – middle graph, p < 0.0001, MW–test). The seizures re– occurred as often as 2.1 ± 0.78 per week when ifenprodil was withdrawn which was not significantly different from the control period (3.87 ± 0.78 per week) (Fig. 6d – middle graph, p = 0.13, MW–test). We did not detect any significant difference in duration between initial (n = 31) and reoccurred (n = 17) seizures (Fig. 6d – right graph, p = 0.86, t–test). Wavelet analysis showed that the average power of analyzed seizures did not vary between control conditions and when ifenprodil administration was stopped as demonstrated by exemplary traces and their spectrograms (Fig. 6e), as well as by spectra of power densities (Fig. 6f –top graph) and their means (Fig. 4f – bottom graph, p = 0.99, 2–way ANOVA). Seizures that were expressed during ifenprodil administration were not included in the analysis as there were insufficient events (n = 2). In summary, the above results demonstrated the strong potential of ifenprodil to suppress chronic seizure in TLE. Ifenprodil–based antagonism of NMDAR_GluN2B_ downregulated chronic seizure occurrence in vivo by ~ 94 %.

## Discussion

The major finding of this study can be summarized as follows: (1) the expression of a specialized subpopulation of NMDAR_GluN2B_ is altered in the TLE hippocampus and corresponds to mossy fiber feedback connectivity in the GC, (2) NMDAR_GluN2B_ is an important component of synaptic transmission at epileptiform GCs, (3) prolonged ifenprodil–mediated antagonism of NMDAR_GluN2B_ during epileptogenesis downregulates the number of seizures during the chronic phase of TLE, (3) inhibition of NMDAR_GluN2B_ function does not affect mossy fiber sprouting formation, (4) NMDAR_GluN2B_ antagonism plays a neuroprotective role in the hippocampus during epileptogenesis, (5) ifenprodil reduces the probability of seizure occurrence during ictogenesis and is effective in chronic seizure suppression in TLE.

### GluN2B in mossy fiber sprouting

To date, three main families of NMDAR subunits have been identified, namely, GluN1, a family of GluN2 subunits (GluN2A–2D), and a pair of GluN3 subunits (GluN3A–GluN3B) (Fan et al., 2014). Native NMDARs are heterotetrameric glutamate–gated channels composed of two obligatory GluN1 subunits and varying expression of a family of GluN2, and, less commonly, also of GluN3 (Fan et al., 2014). Therefore, physiological properties of the NMDAR complex, as well as its coupling to distinct intracellular signaling cascades mostly depends on a unique combination of GluN subunits (Cull–Candy and Leszkiewicz, 2004), which opens the possibility for differential regulation of synapse function in physiology and pathology by particular classes of NMDARs. Our study suggests the GluN2B association with sprouted mossy fiber terminals in animal models of TLE *in vivo*. We show that aberrant synapses formed by the sprouting of mossy fibers may contain a NMDAR_GluN2B_ component. Observations indicate altered expression of NMDAR_GluN2B_ in the DG of TLE mice which morphologically resembles the histological pattern of sprouted mossy fiber terminals.

### Altered GluN2B–dependent EPSC in epileptiform DG

Expression of the GluN2B is spatially and temporarily regulated in the healthy and dysfunctional hippocampus (Akazawa et al., 1994; Monyer et al., 1994). In that regard, the significance of NMDAR_GluN2B_ for TLE–affected hippocampus is not well understood. We show that at dentate GCs of both control and TLE mice, NMDAR_GluN2B_ receptors are activated by the quantal release of glutamate, as demonstrated by the presence of pharmacologically isolated NMDAR_GluN2B_– mediated sEPSCs. Moreover, our analysis shows that the overall contribution of NMDAR_GluN2B_ to the total NMDAR–mediated synaptic transmission is greater in TLE GCs than in control neurons. Therefore our data demonstrate a profound change in the nature of glutamatergic synaptic transmission in epileptiform DG with important participation of NMDAR_GluN2B_ receptors, which might be significant for the pathogenesis of TLE.

### The influence of GluN2B inhibition on epileptogenesis

The role of NMDAR in epileptogenesis has been previously investigated in animal models of TLE. For instance, experiments with dizocyplin (a noncompetitive antagonist of the NMDAR) suggest that administration of an NMDAR antagonist during seizure kindling by stimulation of performant path delays the process of kindling and formation of mossy fiber sprouting in the hippocampus (Sutula et al., 1996). In addition, the general blocking of NMDAR activity in a rat pilocarpine model just before proconvulsant administration eliminates the occurrence of electrographic seizure in EEG in the chronic phase, whereas control animals develop convulsions with typical electrographic seizure (Rice and DeLorenzo, 1998). Therefore, NMDAR function could comprise an ideal target for the prevention of epilepsy development. Unfortunately, since NMDAR play an essential role in the physiology of the brain, antagonists of NMDAR activity, such as dizocyplin, are excluded from medication due to the risk of serious side effects, (Kalia et al., 2008; Kemp and McKernan, 2002). Our data suggest that the NMDAR_GluN2B_ class of receptors play an important role in epileptogenesis. Importantly, NMDAR_GluN2B_ constitutes a very confined target in comparison to global NMDAR population. We show here that selective antagonism of NMDA_GluN2B_ comprises a promising way to affect epileptogenesis and to minimize deleterious side effects of general NMDAR inhibition. As demonstrated with chronic application of ifenprodil in the kainate model of TLE, NMDAR_GluN2B_ antagonism during the first two weeks of epileptogenesis effectively reduces (almost by half) the number of seizures in the chronic phase of the condition. Taken together, our results suggest that ifenprodil interferes with epilepsy development, which may be used to reduce chronic seizure severity in TLE.

### Impact of GluN2B on mossy fiber sprouting formation

Despite the ambiguous role sprouting plays in epileptogenesis, data increasingly confirm that this phenomenon strongly contributes to the development of epilepsy stimulating GCs under conditions of increased excitability or disinhibition of the hippocampal network (Sutula and Dudek, 2007). The experiments carried out *ex vivo* in organotypic cultures have shown that a selective antagonism of the NMDA_GluN2B_ receptor with ifenprodil reduces both mossy fiber sprouting and epileptic discharges of dentate GC (Wang and Bausch, 2004). However, a single injection of ifenprodil after the *status epilepticus* has not been found potent enough to affect mossy fiber sprouting *in vivo* (Chen et al., 2007).

In searching for a mechanism that underlies reduction of chronic seizure frequency upon NMDR_GluN2B_ inhibition during epileptogenesis, we first tested the influence of GC–specific GluN2B downregulation via viral gene transfer on sprouting formation during TLE development and did not find any significant effect. Although the ~30% downregulation of GluN2B we achieved may not be sufficient to affect mossy fiber sprouting, we then systemically inhibited NMDR_GluN2B_ function with ifenprodil for the same purpose. The chosen dose and administration mode of ifenprodil facilitates a potent neuromodulatory effect as demonstrated in further experiments. Despite this fact, we did not find any significant effect of inhibition on sprouting development during epileptogenesis. Our results are in contrast to the effects of NMDR_GluN2B_ observed in organotypic hippocampal cultures (Wang and Bausch, 2004), which could be explained by different physiology of the DG network in the organotypic hippocampal cultures.

### Neuroprotective role of NMDAR_GluN2B_ antagonism during epileptogenesis

A direct link between the neurodegeneration and the etiology of recurrent seizures has not been demonstrated yet (Ono and Galanopoulou, 2012; Bertoglio et al., 2017). While it remains possible that the neurodegeneration observed in TLE patients and animal models of TLE is coincidental, cumulating clinical and experimental data indicate that neurodegenerative conditions and epileptogenesis share a common mechanism (Estrada–Sánchez et al., 2017; Adan et al., 2021).

One of the characteristic features of kainate–induced epilepsy is a profound neurodegenertion in the CA3 region. Due to the fact that kainate receptors are the most abundant in the CA3 region of the hippocampus (Mulle et al., 1998; Carta et al., 2014; Crépel and Mulle, 2015), pyramidal neurons of CA3 are very sensitive to excitotoxic seizure insult (Jarrard, 2002; Wang et al., 2005). CA3 neurodegeneration is also a clinical hallmark of TLE in human patients (Malmgren and Thom, 2012). Here we demonstrated that targeting kainate–induced epileptogenesis with ifenprodil protects the CA3 region against excitotoxic degeneration. Our finding is in line with another study in which 3–day ifenprodil treatment of rats, first subjected to self–sustaining status epilepticus, elicited considerable neuroprotection in CA3 in the chronic phase of the condition (Frasca et al., 2011). We assume that the neuroprotective effects of ifenprodil during epileptogenesis might, to some extent, be responsible for the reduced seizure frequency that we observed in the chronic phase of TLE. However, further studies are required to fully understand the link between these two phenomena.

### The influence of GluN2B inhibition on ictogenesis ex vivo and in vivo

The role of NMDA_GluN2B_ in acute seizure (induced epileptiform discharges in healthy animals) is often confused with the function of NMDA_GluN2B_ in etiology of recurrent seizures in TLE. The latter has not been sufficiently established so far. Therefore, we tested the hypothesis that a selective NMDAR_GluN2B_ block exerts an inhibitory effect on onset of recurrent seizures in the kainate model of TLE. The epileptiform activity can be successfully induced in brain sections from TLE animals under a number of different conditions such as pharmacological disinhibition of the network, adjusted ionic equilibrium of the extracellular solution, and enhanced exposure of the specimen to the oxygen. We provoked ictal–like events in hippocampal brain sections, which reliably reproduced the electrographic seizures recorded *in vivo*. Our data provide evidence that the presence of ifenprodil decreases the likelihood of interictal neuronal network transition toward a seizure, as demonstrated by comparison of the number of ictal–like events in control conditions and when ifenprodil was present. This implicates NMDAR_GluN2B_ as an important factor predisposing neuronal network to seizures and facilitating seizures.

Further, we confirmed this observation *in vivo* by systemic administration of ifenprodil in TLE animlas subjected to chronic vEEG recording. Most treated subjects were completely seizure–free, even though they experienced regular tonic–clonic seizures before the drug administration. The seizure–free condition, however, did not persist after ifenprodil treatment was discontinued.

### Use of ifenprodil to target epileptogenesis and ictogenesis

Ifenprodil (Cerocral©, Furezanil©, Iburonol©, Linbulane©, Technis©, Vadilex©, Vasculodil©) has been approved in some countries for human medication as a competent vasodilator. Here we used ifenprodil to test its potential inhibitory effect on the development of epilepsy and to control recurrent seizures (Radzik et al., 2015).

The potentially preventive effect of ifenprodil against TLE has been already tested in combination with NBQX (Schidlitzki et al., 2017) in the mouse kainate model, but in this context the drug has so far been tested individually. For the first time we report here that prolonged treatment with ifenprodil during epileptogenesis is a promising way to limit the frequency of recurrent seizures in a mouse TLE model.

Ifenprodil has been already proved to be moderately effective in suppressing seizures in models of acute seizure. For instance, Balosso et. al. (2008) described a combined kainic acid and IL1β–provoked convulsions in which animals were treated with ifenprodil prior to induction of seizures. In that approach ifenprodil was shown to reduce the number and duration of seizures, caused by IL1β and kainate, nevertheless, it did not affect seizures when kainate was used alone. Moreover, Yen W et al. (2004) described the use of ifenprodil in an electrical impulse–induced convulsion model, in which ifenprodil showed certain anti–seizure activity in 50% of the animals but was substantially less effective than the other agonists tested. The compound has never been used experimentally *in vivo* against recurrent seizures in TLE. Therefore this is the first study to demonstrate an anti–ictogenic effect of ifenprodil, which is distinct from an anti–convilsive effect in acute seizure models.

In summary, the presented potential of ifenprodil for altering the course of epileptogenesis and ictogenesis provides a rationale for clinical studies on ifenprodil to change its indications for treatments as aimed against TLE.

## Acknowledgments

This work was supported by the National Science Center of Poland (grant 2015/19/D/NZ7/02402) and by the Foundation for Polish Science (BRAINCITY agenda MAB/2018/10). Authors wish to thank the Core Facilities of the Nencki Insititute that were involved in this study – the microscopic data acquisition were performed in the Laboratory of Imaging Tissue Structure and Function, and electrophysiology experiments were conducted with help from the Laboratory of Electrophysiology.

## Notes

### Competing Interest Statement

The authors have declared no competing interest.

